# Millisecond spiking units in dispersed mycelial liquid culture MEA recordings absent in dehydration and fungicidal assays

**DOI:** 10.1101/2025.08.12.669623

**Authors:** Davin Browner, Andrew Adamatzky

**Affiliations:** Unconventional Computing Laboratory, UWE, Bristol, U.K.

## Abstract

The extracellular electrophysiology of dispersed mycelial liquid cultures has not previously been investigated in terms of wide bandwidth frequency components and presence of putative spiking units. Here, we identify stereotypical discrete unit mycelial spikes in 150-3000 Hz filtered extracellular electrophysiological recordings of liquid dispersed mycelial cultures. Hard gold coated and custom designed microelectrode arrays with individual electrode radius of 100 *µ*m were used to identify spikes with frequency components between 150-3000 Hz. The dispersed mycelial cultures were estimated to comprise of a total of 177 spiking units across triplicates (T1= 83, T2=44, T3=50). The triplicates had a combined mean trough-to-peak time of 1.58 ± 0.14 ms. Dehydration and H_2_O_2_ based fungicidal assays showed reduction of the spiking units to zero. Differences in power spectral density (PSD) and Fourier transform analysis of the respective traces suggest that physical structures in the dehydration assay were preserved but that the interface between the mycelium and the electrodes was removed. Biofilm formation was not, therefore, required to detect extracellular spikes using the methods described here. The fungicidal assays removed the discrete unit electrophysiological spike resulting in zero spiking units detected, confirming the physiological origin of the extracellular signals. Extracellular electrophysiology can be studied directly in dispersed liquid cultures of mycelium using suitable microelectrode interfaces without complex media or synthetic interventions such as conductive polymers or nanoparticles.

## Introduction

Fungal electrophysiology has been studied using different intracellular [1, 2] and extracellular [3] methods since the 1960s. In intracellular studies, potentials can be measured via voltage or current using patch clamp interfacing with individual sections of hyphae [4]. However, use of protoplast and heavily modified cell walls are current limitations in patch clamp electrophysiology studies of mycelium t[5]. Extracellular studies are subject to a range of noise sources including environmental, electrochemical and passive ion channel activity. As a result, active signaling can be difficult to parse where signal to noise ratios are not optimal and electrophysiology is not clearly defined [6]. Vibrating probes were used in measurement of single endpoint electrophysiology non-simultaneously along the extent of the proximal region of growing hyphae [3]. The reported current densities showed patterns concomitant with polarising electrical fields. However, these fields could not be conclusively linked to growth phenomena [7]. A further limitation of previous extracellular electrophysiology studies is the lack of wide bandwidth and dynamic recording methods (e.g. 0.1-6000 Hz frequency components at sample rates of 20-30 kHz). Differentiation and parsing of these frequency components would greatly aid in improving understanding of putative stereotypical spike waveforms and how different physiological conditions impact spiking patterns.

In this paper, we identify the discrete unit mycelial spike in 150-3000 Hz filtered extracellular electrophysiological recordings of liquid dispersed mycelial cultures. This confirms that these spikes are present in non-biofilm forming cultures with minimal EPS constituents surrounding the hyphae. Hard gold coated and custom designed microelectrode arrays with individual electrode radius of 100 *µ*m were used to identify spikes with frequency components between 150-3000 Hz. These cultures have minimal exopolymeric substances which are thought to improve the fungal-microelectrode interface. However, a link between lack of EPS contents and comparison with dehydrated and/or dead cells has not been studied in terms of its wide-bandwith (150-3000 Hz) electrophysiological dynamics. The recorded dispersed mycelial cultures were estimated to comprise of a total of 177 spiking units following clustering of detected individual spikes (T1= 83, T2=44, T3=50). Dehydration and H_2_O_2_ based fungicidal assays showed reduction of the spiking units to zero. However, differences in power spectral density (PSD) and Fourier transform analysis of the respective spectra suggest that physical structures in the dehydration assay were preserved but the interface between the mycelium and the electrodes was removed. Biofilm formation is not, therefore, required to detect extracellular spikes using the methods described here. Presence of a liquid interface and growth of hyphal networks on the planar microelectrodes was sufficient to observe spikes. The fungicidal assays removed the discrete unit electrophysiologicalspike. Comparison of viable, dehydrated and non-viable cells will aid in structuring mycelial electrophysiology studies, for example in pathogenic strains, as well as in the development of methods for biosensing and biocomputing.

## Methods

### Liquid cultures

*Hericium erinaceus* cultures were sampled from stocks and grown in flasks for 3 days using sterilised malt liquid culture media (1.5 %). Aeration of the growing cultures was achieved using two syringe Filters (13mm diameter and 0.22 *µ*m filter size). Constant magnetic stirring was utilised during growth to avoid formation of exopolymeric substances (EPS). This approach resulted in viable mycelial liquid cultures with minimal EPS material components.

### Micro-electrode arrays (MEAs)

The microelectrode array (MEA) was a custom printed circuit board (PCB) with 10U” hard gold coating. The MEA had 64 channels including a reference electrode. The single ended circular electrodes in the MEA are arranged in a rectangular grid array with each electrode having a radius of 100 *µ*m and a vertical and horizontal spacing of 700 *µ*m. The exposed electrically conductive surface of the electrodes was equivalent to the diameter at 200 *µ*m each. Recordings were conducted in a TC-5910D shielding box (Tescom, South Korea). The shielding box functioned as a Faraday cage with copper mesh lining and also blocked high frequency noise. With RF shielding effectiveness of up to *>* 80 dB across a wide frequency range, the TC-5910D also acted as a Faraday cage as a result of its conductive copper mesh lining, steel exterior and RF-filtered connectors resulting in a fully enclosed and dark recording environment. The shielding box was placed on an anti-vibration table (Adam Equipment, U.K.) to eliminate mechanical vibration artifacts. Low frequency components and line noise were removed by the bandpass (150-3000 Hz) and common average reference procedures.

### Amplification

The Intan RHD2164 (Intan Technologies, U.S.) was used to amplify the signals from the MEA. It is a 64-channel digital electrophysiology interface chip designed for high-density extracellular recordings. It features 64 low-noise amplifiers with an input-referred noise of 2.4 *µ*V RMS and an input impedance of 1 GΩ // 2 pF. The bandwidth is programmable, with a low cutoff frequency ranging from 0.1 Hz to 500 Hz and a high cutoff frequency from 100 Hz to 20 kHz. The chip includes a 16-bit analog-to-digital converter (ADC) that provides simultaneous sampling of all channels at a maximum sampling rate of 30 kS*/*s per channel. Communication is facilitated through an SPI interface. The MEA was connected via custom housing (connection junctions had a maximum length of 150mm) and routed to the amplifier using an electrode adapter board (Intan Technologies, U.S.). The fungal sample, MEA, and Intan headstages were placed inside the shielding box. The acquisition board controller and computer (which generate digital noise and have high-voltage AC power wiring) were located outside the shielding box. The GND terminal on the Intan headstage was connected to the tissue via a ground reference electrode. The zero-ohm R0 resistor was left in place on the RHD headstage so that REF could be shorted to GND. This resulted in a combined REF/GND electrode that was connected to the fungal sample. The fungal sample was electrically isolated from the shielding box. All recordings were conducted in a temperature controlled environment to minimise thermal noise (19^*°*^C). A butyl rubber lid covered the recording chamber to maintain humidity levels of 90-95 %. Photoelectric noise was negligible due to the dark environment of the interior of the shielding box. The shielding box was locked shut during recordings and was only opened for inspection or changing of samples.

### Data acquisition

The Open Ephys Acquisition Board (OpenEphys, Portugal) was used for data acquisition. It is an open-source interface designed for high-channel-count electrophysiology experiments [8]. In our experiments, a single Intan RHD2164 headstage was connected to the acquisition board via SPI cable. This completed the recording setup and allowed for simultaneously sampling at a per-channel raw recording frequency range of 0.1 to 6500 Hz used in experiments.

### Spike sorting

Recordings had a 0.1 - 6500 Hz bandpass filter implemented using the analog filters of the ADC during acquisition. Following acquisition, a digital bandpass filter was used to isolate activity between 150-3000 Hz using a Butterworth second order filter. This range was selected to be intentionally conservative to avoid picking up residual high frequency signals (unlikely but still possible in the shielded setup). A common reference electrode was used to reduce non-physiological noise further by averaging and common mode noise rejection. Further analysis and spike sorting used the spikeinterface library and kilosort4 spike sorting algorithm [9, 10].

### Fungicidal and dehydration assays

The mycelial liquid culture triplicates were transferred into 12 % H_2_O_2_ solution and left for 2 hours. The resulting cultures were then rinsed with deionised water multiple times and resuspended in liquid media matching that of the initial growth conditions for electrophysiological recordings. Confirmation of non-viability was achieved by subsequent transfer to an agar plate and lack of growth over a period of 5 days for each triplicate. As a result 12 % H_2_O_2_ was found to be an effective fungicidal assay for these samples in terms of growth inhibition. Dehydration of the fungal samples was achieved via drying in an oven for 3 hours at 40 ^*°*^C. Using sterile techniques the dried mycelium was transferred into the MEA recording chamber for electrophysiology recordings.

## Results

The detection of discrete unit spikes and these associated features can be facilitated by use of suitably sized microelectrodes (e.g. with radius of 100 *µ*m). In microscopy of the wild-type samples, septate junctions were observed c. every 50-100 *µ*ms. In Basidiomycota the exterior of the hyphae consists of a cell wall and a semi-permeable bilayer cell membrane called the plasma membrane. The plasma membrane and cell wall are implicated in the transmission of intracellular potentials to the extracellular environment. In general, the plasma membrane and cell wall separates different ion concentrations on the inner and outer sides of the membrane. The preservation of resting membrane potentials is achieved actively within the cell through the regulation of ion movements across the membrane, facilitated by selective ion channels and/or pumps [11]. In active rather than passive ion channels the concentrations are also represented in terms of their charges and the resulting electrochemical gradient forms a membrane potential. In theory and in stereotypical conditions, electrophysiological fluctuations result from opening of ion channels due to chemical or electrical stimulation. The corresponding ions move along a transport dependent and localised electrochemical gradient. The resistance of the membrane is lowered resulting in an inward or outward flow of ions. When measured using intracellular methods this is detected as the transmembrane current. The extracellular fluid is conductive and typically exhibits a low resistance that is not nil. The extracellular current results in a small voltage magnitude that can be measured with proximate electrodes localised to the target tissue. Its current is dependent on Ohm’s law (*U* = *R* ∗ *I*) and is dependent on the properties of the tissue, its relevant active and passive transport phenomena and the operation of voltage and chemical gated ion channels. The extracellular signals are smaller than the transmembrane potentials with the magnitude dependent on the distance of the source to the electrode

Extracellular recordings exhibit signal amplitudes that decrease with increasing distance of the electrode from the signal source. A close interface between the electrode and the cell membrane produces a high signal-to-noise ratio. Typically, distances over 100 *µ*m from an idealised single target cell lead to noise from diffusion phenomena and/or the activity of nearby cells. However, this cannot be applied liberally to all biological systems due to differences in mechanical and electrical properties of tissue as well as the dynamic composition of the extracellular matrix of different cells and organisms. Both the transmembrane current and the extracellular potential follow the same time course, with minor differences due to noise, and are roughly equivalent to the first derivative of the transmembrane potential. If the signal is present in the intracellular environment and is of a high enough magnitude then it should be detectable in the proximal region to the target hyphae, provided suitable EPS or other electrically or ionically conductive substance is present. Agar was found to be a poor intermediary of such signals due to the introduction of electrochemical noise and non-physiological electrical potential shifts such as Donnan potentials. As a result, malt liquid culture media (1.5 %) was preferred in cultivation.

Discrete unit spikes can be defined as having transients of milliseconds duration with frequency components above 150 Hz. For instance, visible spikes following bandpass filtering (e.g. 150-3000 Hz). In extracellular recordings with suitable proximal microelectrodes discrete unit activity should appear as rapid troughs followed by a return to the baseline peak. A further peak prior to the trough may be apparent. In the dispersed mycelial liquid cultures, triplicate 1 exhibited a trough time of 6.00 ms, a peak time of 7.43 ms, a trough-to-peak interval of 1.43 ms, and a half-max width of 1.03 ms. Triplicate 2 exhibited a trough time of 6.00 ms, a peak time of 7.77 ms, a trough-to-peak interval of 1.77 ms, and a half-max width of 1.07 ms. Triplicate 3 exhibited a trough time of 8.60 ms, a peak time of 10.13 ms, a trough-to-peak interval of 1.53 ms, and a half-max width of 16.57 ms. The combined statistics for all triplicates were as follows: trough time 6.87 *±* 1.23 ms, peak time 8.44 *±* 1.20 ms, trough-to-peak interval 1.58 *±* 0.14 ms and half-max width 6.22 *±* 7.31 ms.

The inter-spike interval (ISI) was measured based on definition as the time difference between consecutive spikes in a spike train. The formula for the inter-spike interval is given by:

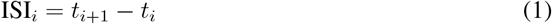

where ISI_*i*_ is the *i*-th inter-spike interval, *t*_*i*_ is the time of the *i*-th spike, and *t*_*i*+1_ is the time of the (*i* + 1)-th spike. Triplicate 1 exhibited a mean inter-spike interval (ISI) of 188.39 ms, a standard deviation of 3421.41 ms, a median ISI of 14.47 ms, and a total of 356,086 ISIs. Triplicate 2 exhibited a mean ISI of 111.07 ms, a standard deviation of 2576.67 ms, a median ISI of 12.47 ms, and a total of 324,773 ISIs. Triplicate 3 exhibited a mean ISI of 229.41 ms, a standard deviation of 2949.76 ms, a median ISI of 18.50 ms, and a total of 180,556 ISIs for the combined total recording time analysed (2700 seconds). The combined inter-spike interval statistics were as follows: mean ISI 167.83 ms, standard deviation 3027.88 ms, median ISI 14.40 ms, and total number of individual ISIs 861,415.

For average spikes per unit (over 900 seconds per triplicate), triplicate 1 exhibited a spikes per unit of 4291.19 *±* 14659.36; triplicate 2 exhibited 7382.20 *±* 19918.00 spikes per unit; and triplicate 3 exhibited 3612.12 *±* 11533.39 spikes per unit. The combined mean was 4867.75 spikes per unit. The mean firing rate across all triplicates was 5.41 *±* 17.18 Hz and the mean amplitudes estimated for the dispersed cultures was 51.04 *±* 63.80*µ*V.

Spike sorting methods used to identify discrete unit extracellular spikes in the dispersed liquid culture triplicates were also applied to the fungicidal and dehydration assays. Parameters were the same as those used in the dispersed liquid culture recordings. In both assays zero spiking clusters were identified for triplicate culture recordings. As a result, we performed power spectral density (PSD) and short-time Fourier transform (STFT) analysis to identify differences in spectra that might explain absences of spiking units in the assays. The dominant frequency for the dispersed liquid culture filtered recording was 175.78 Hz. For the CMR recordings (10-second sample), the mean PSD was −32.80 dB±. As shown comparing raw recordings and STFT plots, shown respectively in Figure 2 (A) and (C), the subthreshold activity can be observed below 100 −150Hz. This activity is related to local field potentials, other physiological sources, isolated growth-related phenomena and non-physiological low-frequency noise. Above these levels the spiking unit activity appeared as higher frequency discrete unit potentials that correspond to the bandpass and common mode reference spike trace in Figure 2 (B). The extensive use of shielding, including high frequency blocking grounded RF box, resulted in high confidence that these discrete unit spikes have an electrophysiological source.

**Figure 1:**
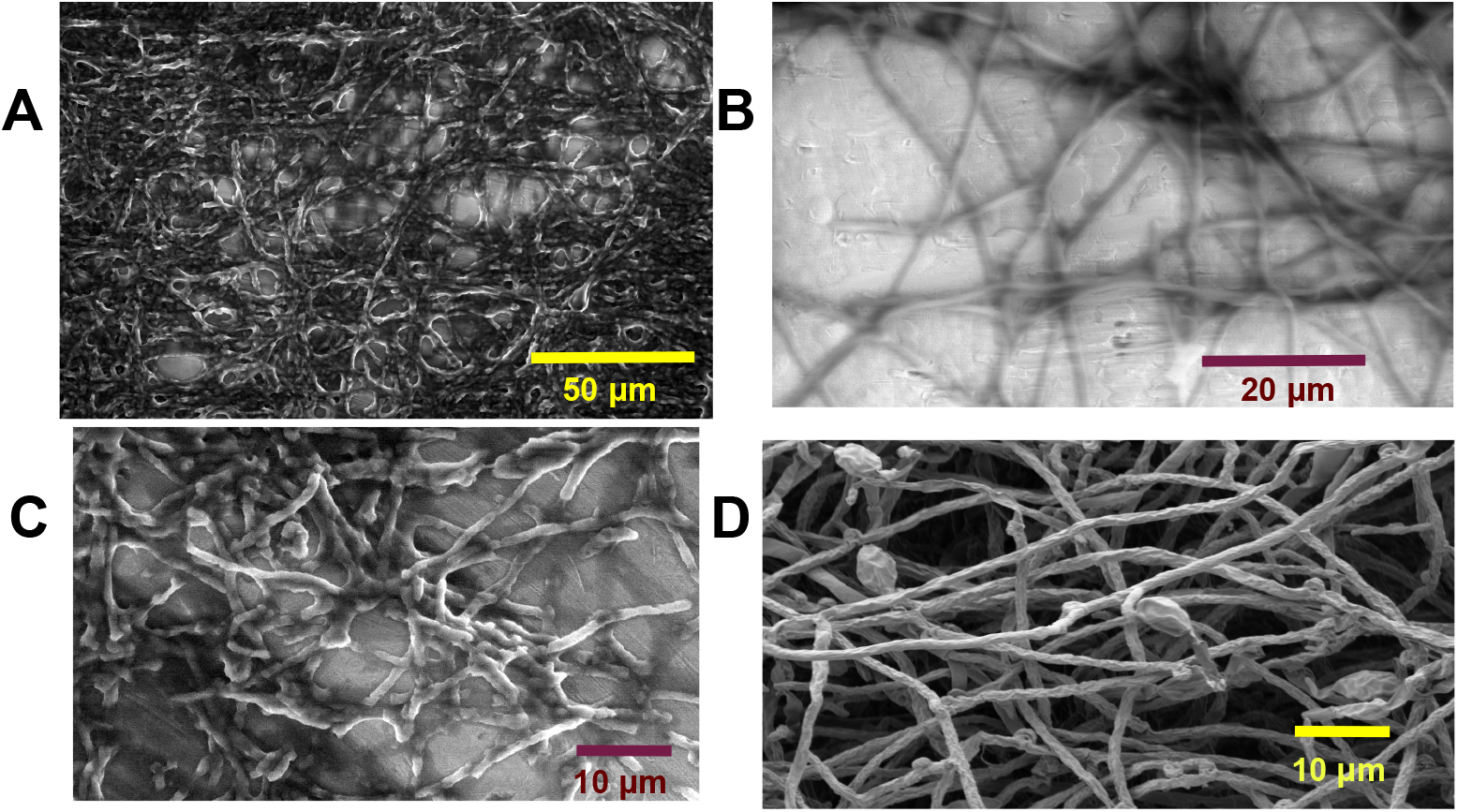
(A) SEM micrograph of wild-type dispersed liquid mycelial sample on SEM stub (HV of 10.0 kV, spot 5.0, magnification of 884x, pressure of 4 Torr and HFW OF 234 *µ*m);(B) Wild-type dispersed liquid mycelial sample growing on the microelectrode surface HV of 10.9 kV, spot 4.0, magnification of 2033x, pressure of 2 Torr and HFW OF 102 *µ*m;(C) Fungicidal assay dispersed liquid mycelial sample on SEM stub HV of 10.0 kV, spot 4.5, magnification of 2467x, pressure of 3 Torr and HFW OF 84 *µ*m;(D) Dehydration assay dispersed liquid mycelial sample on SEM stub HV of 2.0 kV, spot 3.0, magnification of 2352x, pressure of 2.87 ×10^−6^ Torr and HFW OF 88.1 *µ*m.

**Figure 2:**
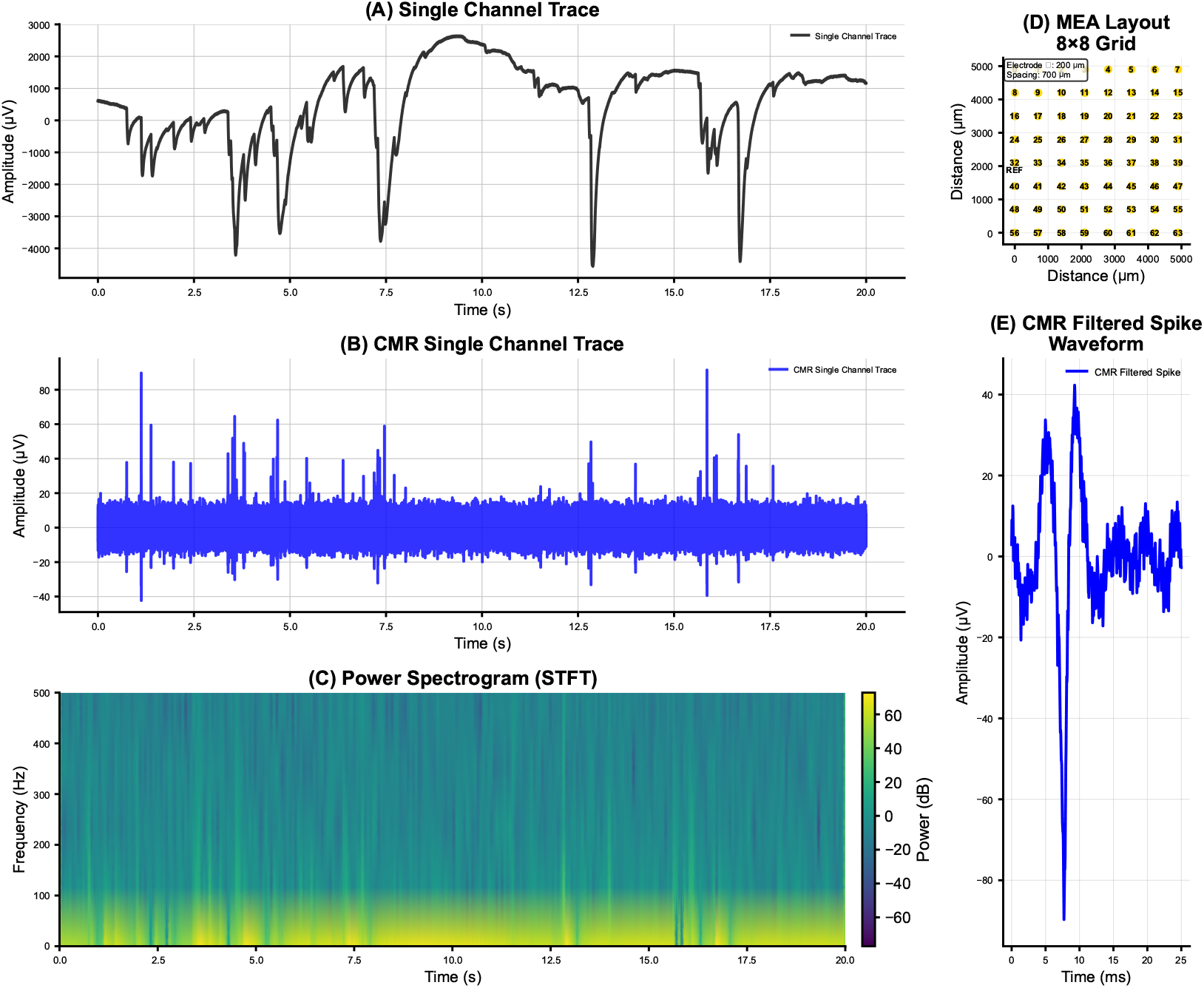
(A) Single channel raw trace (0.1-3000 Hz);(B) Single channel cmr trace (150-3000 Hz);(C) Spectrogram (STFT) showing the discrete unit spikes;(D) 8×8 MEA layout;(E) Isolated discrete unit spike from living dispersed liquid culture mycelium.

**Figure 3:**
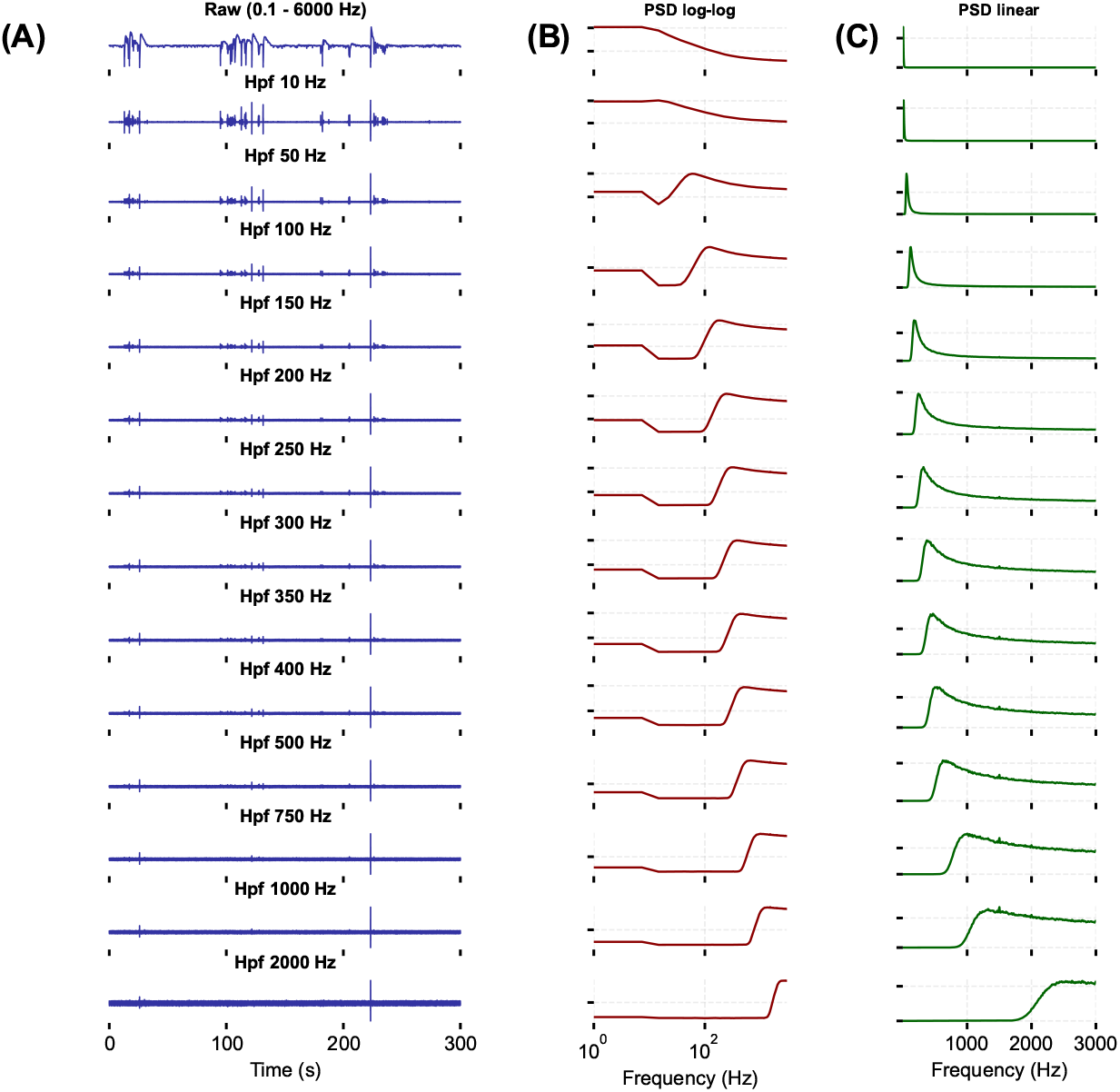
(A) Single channel raw traces at different high pass filters (0.1 - 2000 Hz) with fungal extracellular spikes removed above 2000 Hz;(B) Log-log power spectral density (PSD) at the relevant high pass filters;(C) Linear PSD at the relevant high pass filters.

The interfacial parameters between the dispersed liquid mycelia and the microelectrodes was investigated via a dehydration assay. Zero spiking units were identified across triplicate recordings analysis for dehydrated samples. For the dehydration assay (10-second sample), the mean power spectral density (PSD) was −46.43 dB ± 17.12 dB, and the dominant frequency was 146.48 Hz. No discrete unit activity was observed in the STFT as shown in Figure 5 (C). In addition, subthreshold activity (up to 150Hz) was reduced compared with STFT spectra at this band of the dispersed liquid culture triplicates. However, mean spectra across all channels did not indicate drastically reduced low frequency activity compared with the non-treated dispersed mycelial cultures (outlined in Figure 2). Despite this, comparison of STFT of spiking sections of the recording with random sections of the dehydration assay supports the removal of sub-threshold activity in the dehydration assays. Therefore, an ionic or electrically conductive medium is required for detection of spikes from the extracellular proximal region. In light of these results, we find it unlikely that, given an absence of interfacial conductive components, discrete unit extracellular spikes can be detected at the interface between dehydrated aerial mycelia and electrodes. Use of mycelial cultures in electrophysiological studies and in applications such as sensing or compute may require development of interfacial materials if aerial mycelia are the target structure. An obvious candidate would be agar or other gel-like culture media. However, ion fluxes in agar has been found to produce noise that can be misinterpreted as spikes (e.g. low-frequency Donnan potentials and other similar passive ion fluxes in conductive polymers [12]). Results relating to other gel based conductive interfaces should be interpreted carefully to avoid artifact detection (e.g. Becquerel effect [13]).

The putative physiological origin of spikes in the dispersed liquid culture triplicates was evidenced by the absence of similar spectra in the fungicidal assays. For the H_2_O_2_ fungicidal assay (10-second sample), the mean PSD was −46.51 dB ± 16.97 dB, and the dominant frequency was 498.05 Hz. No discrete unit activity was observed in the STFT as shown in Figure 4 (C). The spectra in combination with the identification of zero spiking units in the spike sorting suggests that the fungicidal assay was successful for all triplicates. This fungicide method resulted in removal of electrophysiological activity and suggests that the electrophysiology observed in the untreated dispersed liquid cultures is physiological in origin and not resulting from electrochemical noise or other sources. Moreover, the spikes are resulting from active cell signaling rather than passive transport of ions such as potentials that may occur in dead but structurally preserved samples.

**Figure 4:**
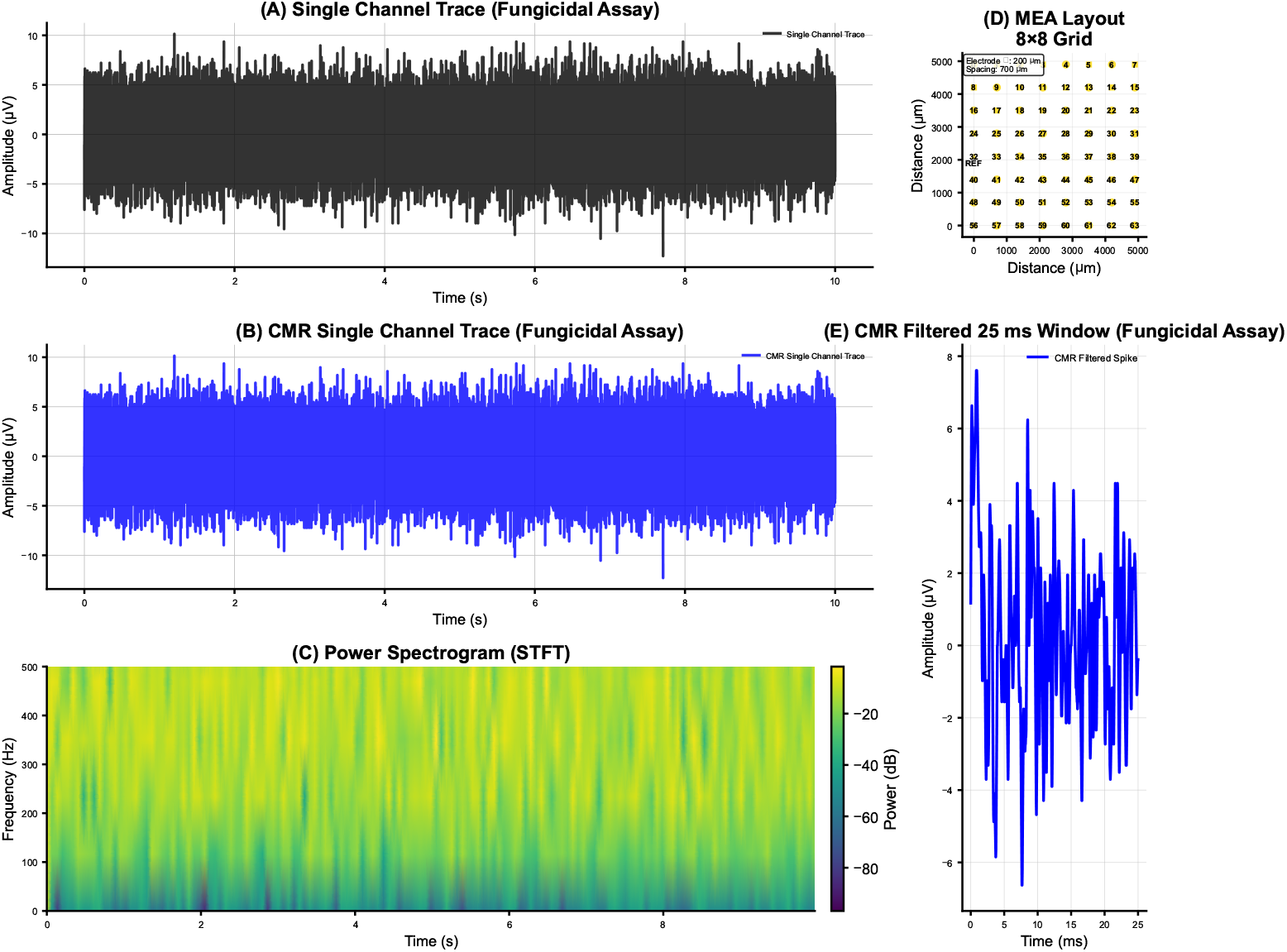
(A) Single channel raw trace (0.1-3000 Hz);(B) Single channel cmr trace (150-3000 Hz);(C) Spectrogram (STFT) showing no evidence of electrophysiological spikes following fungicidal assay;(D) 8×8 MEA layout;(E) 25 ms window shows no evidence of electrophysiological spikes following fungicidal assay.

**Figure 5:**
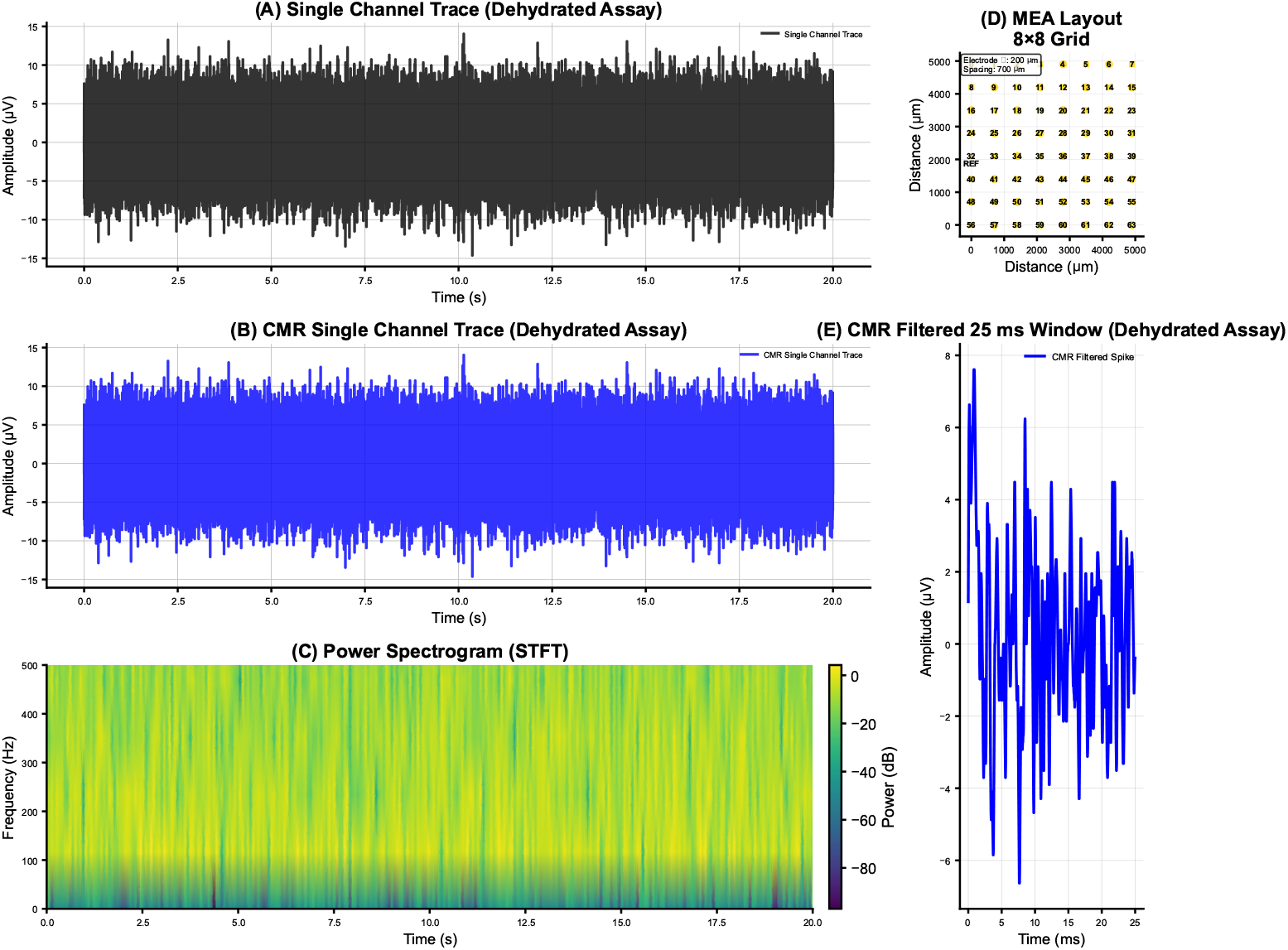
(A) Single channel raw trace (0.1-3000 Hz);(B) Single channel cmr trace (150-3000 Hz);(C) Spectrogram (STFT) showing no evidence of electrophysiological spikes following dehydration assay; (D) 8×8 MEA layout;(E) 25 ms window shows no evidence of electrophysiological spikes following dehydration assay.

**Figure 6:**
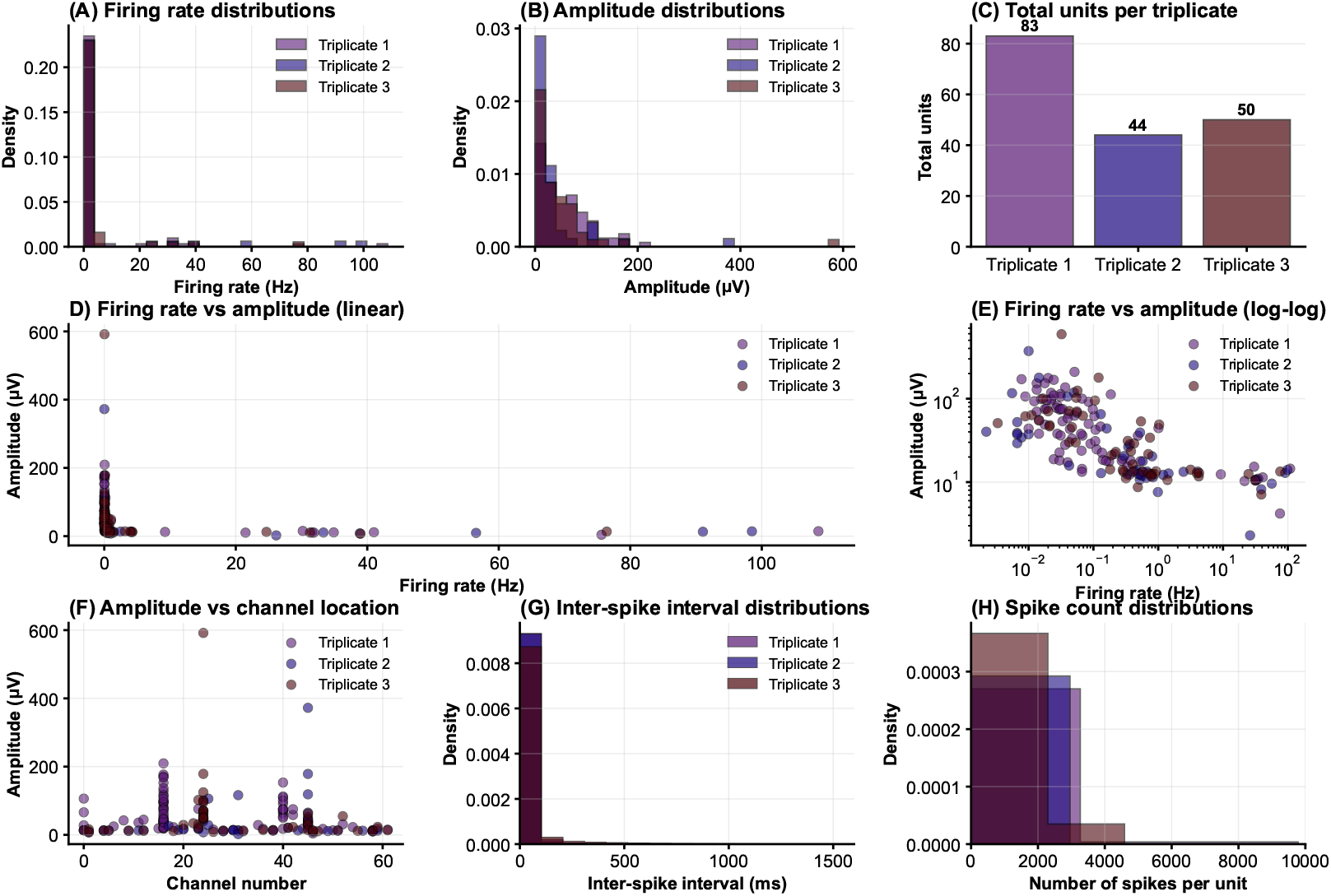
Wild-type *H. erinaceus* discrete unit statistics: (A) Firing rate distribution (Hz);(B) Amplitude distribution (*µ*V); (C) Total units per triplicate (no. units);(D) Firing rate (Hz) vs amplitude (*µ*V); (E) Log-log scale firing rate (Hz) vs amplitude (*µ*V);(F) Amplitude distribution (*µ*V) based on channel location (G) Inter-spike intervals (ms);(H) Spikes per unit.

**Figure 7:**
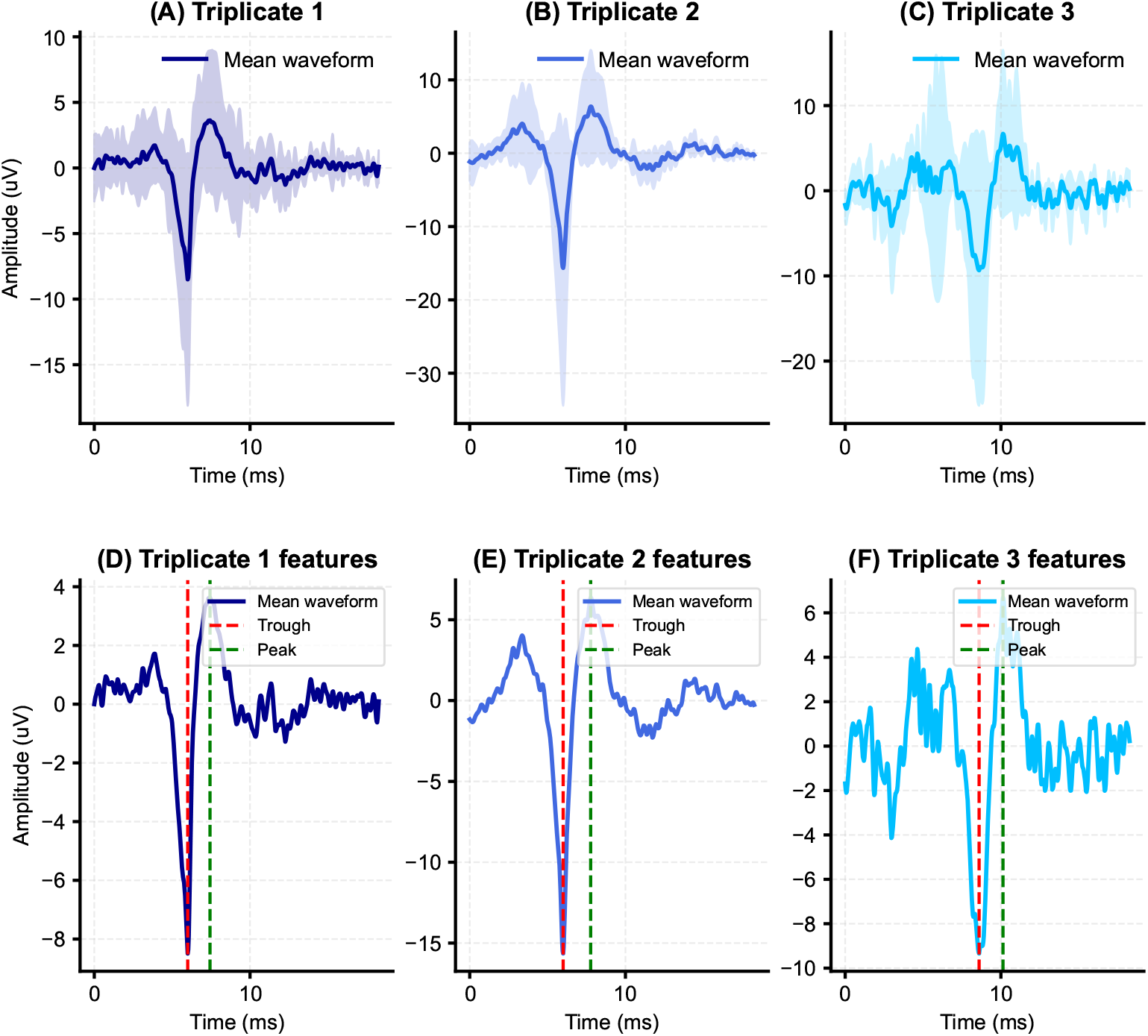
Mean waveforms for the wild-type samples across all units for:(A) Triplicate 1: *H. erinaceus*;(B) Triplicate 2: *H. erinaceus*;(C) Triplicate 3: *H. erinaceus*;(D) Spike waveform properties for triplicate 1;(E) Spike waveform properties for triplicate 2;(F) Spike waveform properties for triplicate 3.

## Discussion

Extracellular electrophysiology can be studied in dispersed liquid cultures of mycelium using a low density microelectrode array. The discrete unit extracellular electrophysiology of mycelium appears to comprise of sub-threshold (≤ 150 Hz) and above threshold discrete unit (≥ 150 Hz) activity. STFT analyses show that the extracellular spikes identified for triplicate recordings have frequency components concomitant with electrophysiological signaling commonly observed in other biological systems (e.g. when neural cultures are recorded using proximal extracellular microelectrodes). A common objection to the presence of putative spiking units in non-neural systems is that the source is electrochemical. However, the frequency components, spike waveforms and regularity of the identified transient signals here does not support an electrochemical origin. Moreover, following dehydration the spikes were absent via removal of the conductive interface between the hyphal spiking unit and the microelectrode. Finally, the spikes were completely removed using fungicidal assays in combination with the liquid nutrient media. Given this combination of factors it is likely that the spikes are physiological in origin.

The combination of Faraday double shielding and RF blocking with a frequency range of 150-3000 Hz for spike detection forms a good foundation for distinguishing between electrophysiological spikes, passive phenomena such as Donnan potentials and environmental noise. Donnan potentials are typically in the 0.1-1 Hz range and therefore are not relevant to the spike detection bandpass utilised here [12]. Similarly mains noise can produce artifacts at 50 Hz. This was below the high pass filter of the bandpass used in sorting and harmonics of this frequency of noise were not detected. All external sensors or sources of noise from the recording chip were configured to the off state (e.g. additional channels or ADC). Grounding and referencing procedures followed guidelines produced by manufacturers [14]. The chamber was sealed from light sources thereby removing the possibility of light-related noise such as the Becquerel effect [13]. Thermal noise artifacts were minimised through the use of a temperature controlled room for all recordings. The recorded potentials are unlikely to be related to physical movement of the mycelium as a result of slow growth dynamics (e.g. electrical artifacts ≤1 Hz).

While the origin of these discrete unit spikes is likely to be physiological there is still the question of what structures produce high frequency spikes. In this sense, the physiological origins of the extracellular spikes are not established. In Basidiomycota, the exterior of the hyphae consists of a cell wall and a semi-permeable bilayer cell membrane called the plasma membrane. In general, hyphae of the lower fungi, i.e. the Glomeromycota, Zygomycota, and Chytridiomycota are sparsely, if at all, septated resulting in a continuous structure [15]. Hyphae of the higher fungi, i.e. the Ascomycota and Basidiomycota, are compartmentalised by septa [15]. These septa contain central pores of up to 500 *nm* that allow streaming of cytoplasm and translocation of organelles like mitochondria and nuclei [15]. Gap junctions in animals and plasmodesmata in plants have pores with a diameter of about 1.5 to 3.0 *nm*. However, the the diameter of the pores of plasmodesmata and gap junctions is dynamic [15]. Heterogenity of the studied hyphae should be considered in extracellular electrophysiology when selecting the spatial dimensions of sorting algorithms as well as microelectrode sizes and layout. In the case where the plasma membrane exhibits a fully continuous structure this would enable cytoplasmic and organelle flow. Without physical impediments or electrically insulating structures propagation of electrical signals would occur in an insulated cable-like structure with less internal electrical resistance [16]. However, hyphal networks punctuated by septal pores at intervals along the extent of individual hypha could have different electrophysiological characteristics when recorded from extracellular environment [16]. In particular, pores could introduce gating junctions as a result of electrically insulating properties or via gating of ion concentrations. The presence of the transmembrane current and resulting extracellular spikes is therefore a complicated proccess to study via one recording method alone.

Early research on fungal electrical signaling suggested the role of proton pumps (H^+^) and various ion channels (Ca^2+^, Cl^-^) [17, 18]. The spontaneous potentials in *Neurospora* were attributed to an electrogenic H^+^ pump or altered membrane selectivity. H^+^ and Cl^-^ were determined to be the primary ions that contribute to the inward currents during potential firing [17]. The spontaneous potential-like firing in *A. bulbosa* and *P. ostreatus* exhibited distinct responses to current injection compared to classical animal models [2]. Later patch clamp studies are of relevance where milliseconds duration spikes in the range of 1-100 ms are visible at approximately 100 times the magnitude of the extracellular spikes observed here post-sorting [19, 20, 5, 21]. It is not entirely surprising that the intracellular potentials recorded from patch clamp and similar methods may have an extracellular correlate given the presence of various relevant ion channels and the transmembrane potential [22]. In theory, the transmembrane current and the extracellular potential follow the same time course, with minor differences due to noise, and are roughly equivalent to the first derivative of the transmembrane potential [2]. Vibrating probe studies are not immediately relevant because of the different recording properties (lock-in amplifiers) and spatial scanning methods [3]. Different ion and voltage-gated channels have been described in filamentous fungi [22]. Voltage-gated proton channels were related to the fungal Hv channels and conformational changes that resulted in gate opening and proton conduction. Fungal Hvs share 20–29 % sequence identity with the human voltage-gated proton channel hHv1 [23]. Whole-genome sequencing has facilitated the identification of genes likely encoding homologues of K^+^, Ca^2+^, transient receptor potential (Trp), and mitochondrial Ca^2+^ uniporter channels [24]. Homology searches for Basidiomycota have shown presence of putative voltage-gated channels in response to multiple ions (including Ca^2+^, Cl^+^, H^+^ and Na^+^) and the signaling molecule glutamate [25]. The combination of homology searches and ion channel isolating patch clamp studies are useful for identification of these putative voltage-gated channels. However, once the channels have been identified the short duration and non-physiological conditions (i.e. protoplast extrusion) of current patch clamp methods are major limitations in detection of discrete unit potential spikes.

## Conclusion

In this paper we outlined methods to record and detect discrete unit spikes in non-biofilm forming *H. erinaceus* liquid cultures using a custom designed hard gold coated microelectrode array. Offline spike sorting of the discrete unit potentials estimated that there were a total of 177 individual spiking units across all wild-type triplicates with a combined mean trough-to-peak time of 1.58 ± 0.14 ms. A dehydration assay confirmed, across triplicates, that a conductive medium is required for signal transmission and detection by proximal microelectrodes. However, the presence of EPS is not required to detect spikes where mycelial constituents are dispersed in the nutrient media. A fungicidal assay utilising 12% w/v H_2_O_2_ solution resulted in zero spiking units confirming physiological and electrophysiological origin of the previously detected spikes. The methods, hardware and signal processing techniques presented in this paper are expected to aid in the development of standardisation of studies of extracellular electrophysiology of mycelium.

## Acknowledgment

The authors would like to thank David Patton for assistance with the environmental scanning electron microscopy. The research has been conducted under the framework of the FUNGATE-RIA (www.fungateria.eu) project, which has received funding from the European Union’s HORIZON-EIC-2021-PATHFINDER CHALLENGES programme under grant agreement No. 101071145. It is co-funded by the UK Research and Innovation grant No. 10048406.

## Data availability

The datasets used and/or analysed during the current study available from the corresponding author on reasonable request.

## References

[1] Clifford L Slayman and Carolyn W Slayman. Measurement of membrane potentials in neurospora. Science, 136(3519):876–877, 1962.

[2] S Olsson and BS Hansson. Action potential-like activity found in fungal mycelia is sensitive to stimulation. Naturwissenschaften, 82(1):30–31, 1995.

[3] Robert F Stump, Kenneth R Robinson, Ruth L Harold, and Franklin M Harold. Endogenous electrical currents in the water mold blastocladiella emersonii during growth and sporulation. Proceedings of the National Academy of Sciences, 77(11):6673–6677, 1980.

[4] Katherine L Perkins. Cell-attached voltage-clamp and current-clamp recording and stimulation techniques in brain slices. Journal of neuroscience methods, 154(1-2):1–18, 2006.

[5] Tanja Pajić, Katarina Stevanović, Nataša Todorović, Steva Lević, Svetlana Savić Šević, Dejan Pantelić, Miroslav Živić, Mihailo D Rabasović, and Aleksandar J Krmpot. Laser nano-surgery of fungal cell wall to enable patch clamping. In European Molecular Imaging Meeting: 18th Annual Meeting of the European Society for Molecular Imaging: EMIM 2023; 2023 Mar 14-17; Saltzburg, Austria, page 1095. European Society for Molecular Imaging, 2023.

[6] Richard Mayne, Nic Roberts, Neil Phillips, Roshan Weerasekera, and Andrew Adamatzky. Propagation of electrical signals by fungi. Biosystems, 229:104933, 2023.

[7] NAR Gow and BM Morris. The electric fungus. Botanical Journal of Scotland, 47(2):263–277, 1995.

[8] Joshua H Siegle, Aarón Cuevas López, Yogi A Patel, Kirill Abramov, Shay Ohayon, and Jakob Voigts. Open ephys: an open-source, plugin-based platform for multichannel electrophysiology. Journal of neural engineering, 14(4):045003, 2017.

[9] Alessio Paolo Buccino, Cole Lincoln Hurwitz, Samuel Garcia, Jeremy Magland, Joshua H Siegle, Roger Hurwitz, and Matthias H Hennig. Spikeinterface, a unified framework for spike sorting. Elife, 9:e61834, 2020.

[10] Marius Pachitariu, Shashwat Sridhar, Jacob Pennington, and Carsen Stringer. Spike sorting with kilosort4. Nature Methods, 21(5):914–921, 2024.

[11] Erwin Neher, Alain Marty, and Elizabeth Fenwick. Ionic channels for signal transmission and propagation. Progress in Brain Research, 58:39–48, 1983.

[12] Hiroyuki Ohshima and Shinpei Ohki. Donnan potential and surface potential of a charged membrane. Biophysical journal, 47(5):673–678, 1985.

[13] PJ Hillson and Eric Keightley Rideal. The becquerel effect in the presence of dyestuffs and the action of light on dyes. Proceedings of the Royal Society of London. Series A. Mathematical and Physical Sciences, 216(1127):458–476, 1953.

[14] Intan Technologies. Noise reduction techniques for intan recording systems. Technical note, Intan Technologies, 2021. Accessed: 2025-04-21.

[15] Arend F van Peer, Wally H Müller, Teun Boekhout, Luis G Lugones, and Han AB Wösten. Cytoplasmic continuity revisited: closure of septa of the filamentous fungus schizophyllum commune in response to environmental conditions. PLoS one, 4(6):e5977, 2009.

[16] TV Potapova and Boitzova LJu. Structure, function, regulation: experimental analysis in groups of non-excitable cells coupled via permeable junctions. Membrane & Cell Biology, 11(6):817–829, 1998.

[17] Clifford L Slayman, W Scott Long, and Dietrich Gradmann. “action potentials” in neurospora crassa, a mycelial fungus. Biochimica et Biophysica Acta (BBA)-Biomembranes, 426(4):732–744, 1976.

[18] Franklin M Harold, Darryl L Kropf, and John H Caldwell. Why do fungi drive electric currents through themselves? Experimental mycology, 9(3):3–86, 1985.

[19] Stephen K Roberts, Graham K Dixon, Stuart J Dunbar, and Dale Sanders. Laser ablation of the cell wall and localized patch clamping of the plasma membrane in the filamentous fungus aspergillus: characterization of an anion-selective efflux channel. The New Phytologist, 137(4):579–585, 1997.

[20] Anne-Alienor Very and Julia M Davies. Laser microsurgery permits fungal plasma membrane single-ion-channel resolution at the hyphal tip. Applied and environmental microbiology, 64(4):1569–1572, 1998.

[21] Natalia N Levina, Roger R Lew, Geoffrey J Hyde, and I Brent Heath. The roles of ca2+ and plasma membrane ion channels in hyphal tip growth of neurospora crassa. Journal of Cell Science, 108(11):3405–3417, 1995.

[22] Gabriella Houdinet, Carmen Guerrero-Galán, Benjamin D Rose, Kevin Garcia, and Sabine D Zimmermann. Secrets of the fungus-specific potassium channel tok family. Trends in Microbiology, 31(5):511–520, 2023.

[23] Chang Zhao and Francesco Tombola. Voltage-gated proton channels from fungi highlight role of peripheral regions in channel activation. Communications Biology, 4(1):261, 2021.

[24] David L Prole and Colin W Taylor. Identification and analysis of cation channel homologues in human pathogenic fungi. 2012.

[25] Xinjiang Cai. Ancient origin of four-domain voltage-gated na+ channels predates the divergence of animals and fungi. The Journal of membrane biology, 245:117–123, 2012.

